# Lung-Targeting Interleukin-10 mRNA Lipid Nanoparticles Ameliorate Acute Lung Injury

**DOI:** 10.64898/2026.01.22.701009

**Authors:** Yuqin Men, David O. Popoola, Yuqi Song, Zhi Cao, Robert Gardner, Rezwana Karim, Chunyan Wang, Nathan Tucker, Robert Cooney, Qinghe Meng, Yamin Li

**Affiliations:** Department of Pharmacology, The State University of New York, Upstate Medical University, Syracuse, NY 13210, USA; Department of Surgery, The State University of New York, Upstate Medical University, Syracuse, NY 13210, USA; Sepsis Interdisciplinary Research Center (SIRC), The State University of New York, Upstate Medical University, Syracuse, NY 13210, USA

**Keywords:** Lung-targeting, lipid nanoparticle, IL-10, mRNA delivery, acute lung injury

## Abstract

Acute respiratory distress syndrome (ARDS) is the most severe manifestation of acute lung injury (ALI), characterized by diffuse pulmonary inflammation, impaired gas exchange, and high morbidity and mortality. Despite its clinical significance, no specific or effective pharmacological therapies are currently available for its treatment. In this study, we developed a lung -targeted mRNA-sulfonium lipid nanoparticle (mRNA/sLNP) delivery system for the treatment of ALI in a mouse model. We first optimized sulfonium lipid structures, and the optimized sLNP was comprehensively characterized and subsequently loaded with interleukin-10 (IL-10) mRNA. In a lipopolysaccharide (LPS)-induced ALI mousemodel, IL-10/sLNP demonstrated both prophylactic and therapeutic efficacy, significantly attenuating pulmonary and systemic inflammation, restoring barrier integrity, and reducing tissue injury.

## Introduction

Acute lung injury (ALI) encompasses a class of severe inflammation conditions in the lungs caused by direct or indirect injuries.^1-3^ These conditions are characterized by an uncontrolled inflammatory response, including the release of pro-inflammatory cytokines, neutrophil infiltration, damage to the alveolar-capillary barrier, and pulmonary edema. A more severe form of ALI, acute respiratory distress syndrome (ARDS), commonly affects critically ill patients, including those with COVID-19.^4-6^ Current treatment strategies for ALI/ARDS primarily rely on protective mechanical ventilation and supportive critical care.^7-9^ Despite advances in critical care management, ALI/ARDS remains a leading cause of mortality.^10-12^ The high mortality rates underscore the urgent need for effective therapeutic interventions. Corticosteroids such as dexamethasone are widely used for treating adult patients with ARDS. However, systemic corticosteroid use is usually accompanied by various adverse effects.^13, 14^ Alternative pharmacological approaches have been explored, such as neuromuscular blockers, inhaled nitric oxide, antioxidants, surfactant therapy, and anticoagulation. ^15-19^ Despite all the promising advances, safe and effective pharmacological treatment for ALI/ARDS remains elusive.

Interleukin-10 (IL-10) is a pleiotropic cytokine with a critical role in modulating inflammation and maintaining immunological homeostasis.^20, 21^ Its signaling cascade is primarily mediated by the Jak1/Tyk2/STAT3 pathway, leading to inhibition of MAPK activation, NF-κB nuclear translocation, and pro-inflammatory gene expression.^22-24^ IL-10 therapy has demonstrated potent immunomodulatory effects in various inflammatory diseases, such as rheumatoid arthritis, asthma, inflammatory bowel disease, and psoriasis.^25-29^ Importantly, IL-10 inhibits LPS- and bacteria-mediated induction of pro-inflammatory cytokines such as TNF-α, IL-1β, IL-12, and IFN-γ in TLR-activated myeloid cells, and it has shown therapeutic benefits in ALI and endotoxemia.^30-32^ However, due to its suboptimal pharmacokinetics and non-specific biodistribution profile, the therapeutic efficacy of systemic administration of IL-10 cytokine has been limited.

In this study, we developed a lung-targeting sulfonium lipid nanoparticle (sLNP) system for IL-10 mRNA delivery as a potential treatment for ALI/ARDS. Our group previously reported the design and synthesis of sulfonium lipids and demonstrated their ability to efficiently deliver mRNA to the lungs in mice.^33^ Sulfonium lipids are a type of cationic lipid material, chemically different from amine-based cationic lipids.^34-36^ In this study, we investigated how variations in the head and tail structures of sulfonium lipids influence biodistribution and lung-targeting efficiency. We also examined the correlation between the physicochemical properties of sLNPs and their mRNA delivery performance in the lungs. We delivered IL-10 mRNA and evaluated its prophylactic and therapeutic efficacy in an LPS-induced ALI mouse model. A schematic illustration of this study is shown in **Figure 1A**.

**Figure 1.**
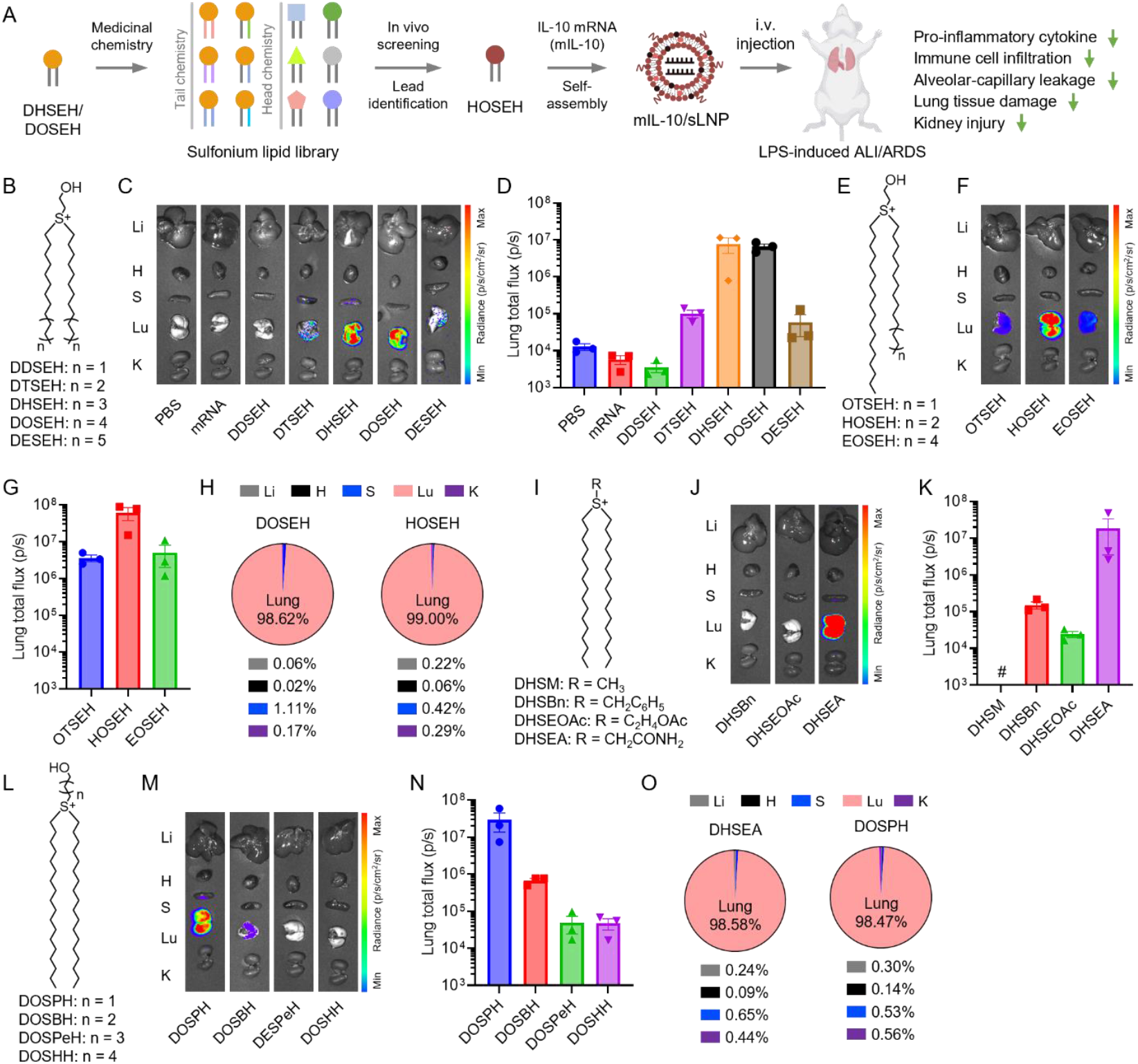
Sulfonium lipid nanoparticle (sLNP)-enabled systemic delivery of mRNA to the lungs for the treatment of acute lung injury. (A) Schematic illustration of the sulfonium lipid structure optimization using a medicinal chemistry approach, and sLNP-mediated, lung-specific, IL-10 mRNA delivery in mice. (B, E, I, and L) Chemical structures of sulfonium lipids with various head and tail structures. (C, F, J, and M) Representative ex vivo bioluminescence images of mouse organs treated with mFLuc/sLNP formulations. (D, G, K and N) Quantification of total bioluminescence flux from lungs. Data are presented as mean ± SEM. N = 3. (H and O) Percentage of background-subtracted bioluminescence signal originating from each organ.

### Results and discussion

#### Effects of sulfonium lipid tail structure on lung-targeted mRNA delivery

Building on our previously reported hydroxyethyl-headed DHSEH and DOSEH lipids, we synthesized several groups of new sulfonium lipids with varying head, linker, and tail structures.^33, 37, 38^ We first examined a group of sulfonium lipids with varying hydrophobic tail lengths, ranging from dodecyl (C12) to eicosyl (C20), specifically DDSEH, DTSEH, and DESEH (**Figure 1B**). Four-component sLNP formulations were prepared via self-assembly using sulfonium lipids, cholesterol, phospholipid DOPC, and PEG-lipid PEG2k-DMG.^33^ Firefly luciferase mRNA (mFLuc) was encapsulated in the sLNPs, and the resulting formulations were administered to adult C57BL/6 mice via intravenous injection. Expression of mRNA-encoded FLuc protein in major organs (i.e., liver, heart, spleen, lung, and kidneys) was assessed 6 h post-injection using *ex vivo* bioluminescence imaging (**Figure 1C**). As expected, similar to the PBS control group, no bioluminescence signal was detected in the organs of mice injected with free mRNA, likely due to its inherent instability and inability to penetrate biological membranes. Consistent with our previous findings, DHSEH and DOSEH, which feature hexadecyl (C16) and octadecyl (C18) tails, respectively, induced strong bioluminescence signals, predominantly in the lungs.^33^ Bioluminescence was also observed in the lungs of mice treated with DTSEH and DESEH, whereas no signal was detected in the DDSEH group.

Quantification of total lung bioluminescence flux revealed that DHSEH (7.89 × 10^6^ p/s) and DOSEH (6.61 × 10^6^ p/s) achieved the highest intensities, while DTSEH (1.01 × 10^5^ p/s) and DESEH (5.90 × 10^4^ p/s) exhibited moderate activity (**Figure 1D**). In contrast, DDSEH (3.52 × 10^3^ p/s) was inactive, with flux levels comparable to the PBS (1.27 × 10^4^ p/s) and free mRNA (5.74 × 10^3^ p/s) control groups. These findings indicate that the optimal linear alkyl tail length for hydroxyl ethyl-headed sulfonium lipids ranges from C16 to C18. Shorter or longer tail lengths resulted in diminished lung-targeted mRNA delivery in mice. Previous studies suggest that lipid tail length plays a crucial role in determining both delivery efficiency and cell/tissue selectivity *in vitro* and *in vivo*. While no widely-accepted rationale exists to explain this phenomenon, it is likely that variation in the tail length may influence multiple factors, including LNP formation, colloidal stability, and nano-bio interactions.^39-41^

Next, we synthesized and evaluated a small group of sulfonium lipids with asymmetric tail structures (**Figure 1E**). Incorporating asymmetry into lipid molecule design has emerged as a promising strategy to improve delivery. It has been shown that combinatorial lipids libraries synthesized via multi-component reactions often feature asymmetric tails, and these lipids exhibited enhanced efficacy and cell-type specificity.^42, 43^ Here, sulfonium lipids containing an octadecyl tail paired with either a tetradecyl (OTSEH), hexadecyl (HOSEH), or eicosyl (EOSEH) tail were formulated into sLNPs and loaded with mFLuc. Following intravenous injection in adult mice, bioluminescence signals were detected primarily in the lungs across all groups (**Figure 1F**). Among these formulations, HOSEH demonstrated the highest efficacy, with a total lung flux of 6.12 × 10^7^ p/s (**Figure 1G**), outperforming both OTSEH (3.62 × 10^6^ p/s) and EOSEH (5.03 × 10^6^ p/s). Notably, HOSEH achieved an 8-to 9-fold increase in lung bioluminescence compared to DHSEH and DOSEH, highlighting its potency. OTSEH and EOSEH also exhibited superior delivery efficacy relative to their counterparts with symmetric tails, DTSEH and DESEH. Lipid species with asymmetric tails are abundant in biological membranes, yet the impact of tail asymmetry on membrane properties, particularly membrane dynamics, remains poorly understood.

Koynova et al. reported that ethylphosphatidylcholine lipids with asymmetric chains enhance lamellar-to-nonlamellar phase transitions, which may contribute to membrane fusion and cargo release.^44^ Meka et al. reported that ammonium cationic lipids bearing asymmetric tails enhanced membrane fusion and endosomal escape, leading to improved transfection.^45^ Prior studies suggest that tail chain asymmetry may affect membrane dynamics across molecular to mesoscopic scales, while the underlying mechanism of asymmetry-induced delivery enhancement in sLNP merits further investigation.

To further investigate lung specificity, we analyzed the bioluminescence distribution of the two top-performing sLNPs: DOSEH and HOSEH (**Figure 1H**). Quantification analysis revealed that 98.6% and 99.0% of the total bioluminescence signal originated from the lungs of DOSEH and HOSEH-treated mice, respectively. Minimal signal (<1.2%) was detected in other organs (i.e., liver, heart, spleen, and kidneys), underscoring the exceptional lung-targeting specificity of these formulations. The specificity of sLNP-mediated mRNA delivery is expected to be advantageous in minimizing off-target mRNA delivery and potential systemic adverse effects in future translational studies. It is noteworthy that the hydroxyethyl-headed lipids examined in this study displayed either no activity or, when active, yielded luciferase expression that was highly restricted to lung tissue. In a previous study, we observed that incorporating asymmetrical and branched tail structure (i.e. 2-hexyl-decyl) into sulfonium lipids redirected mRNA/sLNP expression to both the spleen and lungs (with comparable bioluminescence flux) following intravenous injection in adult mice.^37^ Although the underlying mechanism remains unclear, this result indicates that modifying the tail architecture of sulfonium lipids may serve as a viable strategy to reprogram sLNP organ tropism beyond the lungs.

#### Effects of sulfonium lipid head on lung-targeted mRNA delivery

Next, we examined the effects of sulfonium lipid head groups on sLNP-mediated mRNA delivery. A series of sulfonium lipids was synthesized by replacing the hydroxyethyl group in DHSEH (**Figure 1I**) with methyl (DHSM), benzyl (DHSBn), ethyl acetate (DHSEOAc), and acetyl amide (DHSEA). These lipids were formulated into mFLuc-loaded sLNPs and administered to mice via intravenous injection (**Figure 1J**). During formulation, DHSM-based sLNPs were found to be unstable, exhibiting precipitation following self-assembly, and were therefore excluded from further evaluation. DHSEA (1.84 × 10^7^ p/s) exhibited the strongest bioluminescence signal in lung tissues, comparable to DHSEH and DOSEH (**Figure 1K**). DHSBn (1.51 × 10^5^ p/s) induced moderate bioluminescence in the lungs, similar to DTSEH and DESEH. No signal was detected in the DHSEOAc group (2.43 × 10^4^ p/s), where the total flux was indistinguishable from the PBS control. These results indicate that head group substitution significantly impacts mRNA delivery. The hydroxyl group in DHSEH is critical, as replacing it with methyl group increased hydrophobicity and destabilized the self-assembly structures, while caging it with an acetyl ester group (DHSEOAc) resulted in a complete loss of activity. Benzyl substitution reduced delivery efficacy, whereas acetyl amide substitution neither enhanced nor diminished delivery efficiency.

Additionally, we synthesized a group of DOSEH analogs featuring different linker lengths between the hydroxyl and sulfonium groups (**Figure 1L**): DOSPH (propyl, C3), DOSBH (butyl, C4), DOSPeH (pentyl, C5), and DOSHH (hexyl, C6). DOSPH (2.93 × 10^7^ p/s) and DOSBH (6.77 × 10^5^ p/s) showed strong bioluminescence in the lungs (**Figure 1M**), while negligible signals were observed in mice treated with DOSPeH (4.87 × 10^4^ p/s) or DOSHH (4.69 × 10^4^ p/s), which was further confirmed by the total lung flux quantification (**Figure 1N**). These findings indicate that the linker length between the hydroxyl and sulfonium groups is critical for mRNA delivery. Carbon spacers of two (DOSEH) and three (DOSPH) carbons resulted in specific and effective lung - targeted mRNA delivery, while longer spacers gradually reduced delivery efficacy.

Lastly, we quantified tissue specificity for the top-performing sLNPs (**Figure 1O**). Greater than 98% of bioluminescence signals originated from the lungs in both DHSEA- and DOSPH-treated mice, highlighting the exceptional lung specificity of these sLNP formulations. Our results demonstrate that hydroxyl- and amide-substituted sulfonium lipids with certain linker lengths are effective for mRNA delivery, whereas methyl-, benzyl-, and acetyl ester-substituted lipids are either unstable or less effective.

#### Correlation of sLNP properties with lung-targeted mRNA delivery

Building upon the successful optimization of sulfonium lipid structures, we further investigated the structure-activity relationship of sulfonium lipids by examining how sLNP physicochemical properties correlate with *in vivo* delivery efficacy. To this end, we selected three top-performing (DOSEH, DOSPH, and HOSEH), three moderately performing (DHSBn, DOSBH, and OTSEH), and three low-performing (DDSEH, DHSEOAc, and DOSPeH) sLNPs for comprehensive characterization. We first assessed the impact of sulfonium lipid hydrophobicity on mRNA delivery. The calculated LogP (cLogP) values for the nine lipids ranged from 8.16 to 10.12, resembling the hydrophobicity of phospholipids (**Figure 2A**). DDSEH, with the shortest hydrophobic tail (C12), was the least hydrophobic (cLogP = 8.16), while DOSPeH, with the longest hydrophobic tail (C18) and linker length (C5) in the nine lipids, was the most hydrophobic (cLogP = 10.12). However, no significant correlation was observed between cLogP and lung-targeted mRNA delivery (Spearman correlation coefficient, r = 0.2, p = 0.6134; **Figure 2B**). At the supramolecular level, the sLNPs exhibited an average hydrodynamic size of 100–200 nm (**Figure 2C**), a polydispersity index (PDI) of 0.1–0.3, a zeta potential of 43–55 mV, and an mRNA encapsulation efficiency of 94–99% (**Figure 2G**). Spearman correlation analysis revealed no significant relationship between these parameters and lung mRNA delivery efficacy, with correlation coefficients ranging from -0.4167 to 0.0667 (p > 0.05; **Figures 2D, E, F**, and **H**). Collectively, these results suggest that neither molecular hydrophobicity (cLogP) nor supramolecular characteristics (size, PDI, zeta potential, and encapsulation efficiency) are directly correlated with sLNP-mediated *in vivo* lung-targeted mRNA delivery.

**Figure 2.**
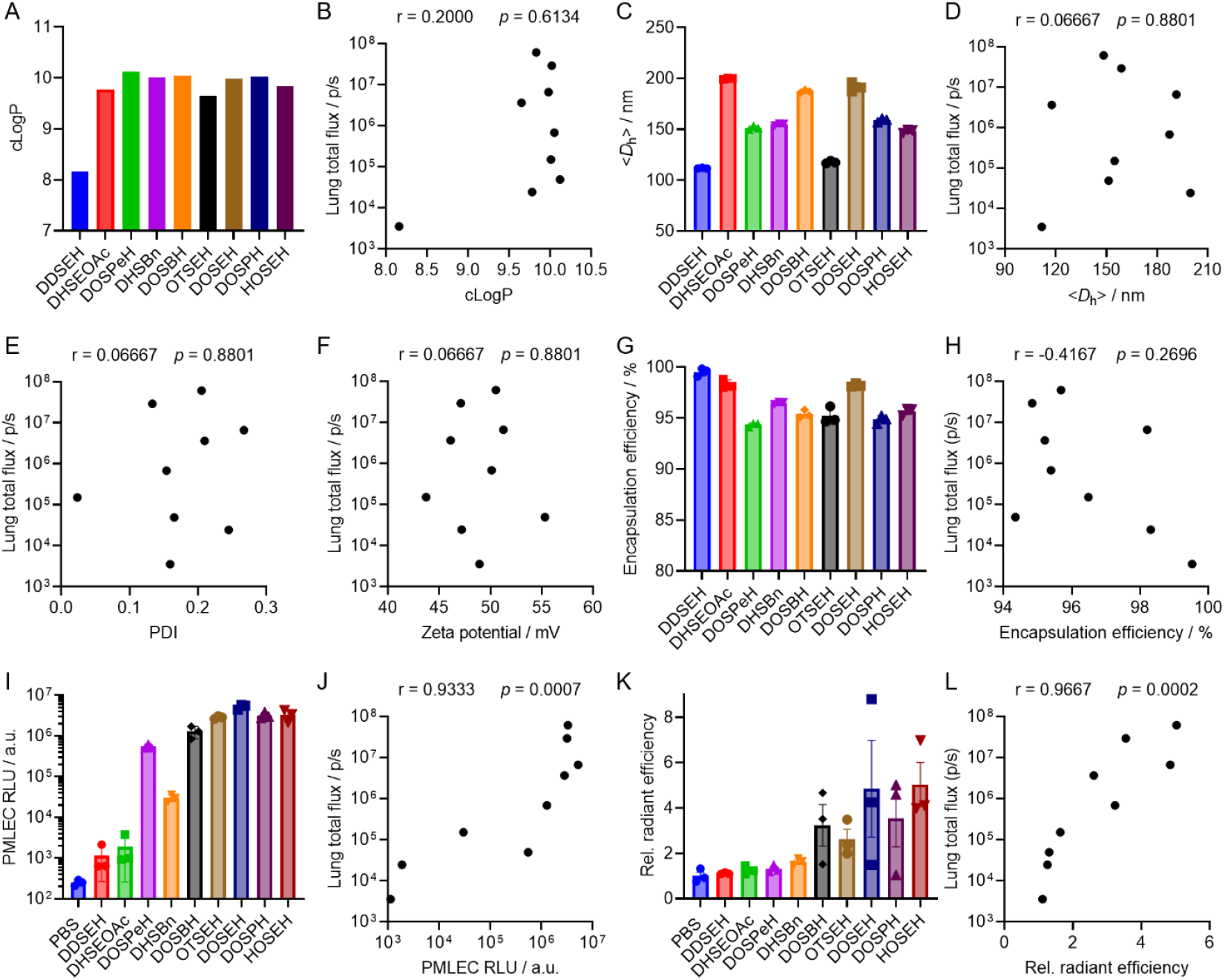
Structure-activity relationship study of sLNP formulations. (A) cLogP of sulfonium lipids, (C) <*D*_h_>, (G) mRNA encapsulation efficiency, (I) in vitro PMLEC transfection efficiency, and (K) in vivo lung accumulation efficiency of sLNP formulations. Spearman correlation analysis of in vivo mRNA delivery efficacy to lungs and (B) cLogP, (D) <*D*_h_>, (E) PDI, (F) zeta potential, (H) encapsulation efficiency, (J) PMLEC transfection efficiency, and (L) in vivo lung accumulation efficiency. Data are presented as mean ± SEM. N = 3.

Next, we investigated whether the transfection efficiency observed *in vitro* correlates with the *in vivo* lung-targeted mRNA delivery efficiency. Since no correlation was observed between the physicochemical properties of sLNPs and lung-specific delivery, cell-free analyses appeared less informative in this context. In our previous work, we identified lung endothelial cells as the primary targets of DOSEH-mediated mRNA delivery and protein expression. Therefore, we selected primary mouse lung endothelial cells (PMLECs), isolated using magnetic bead separation, to assess FLuc mRNA transfection efficiency in vitro. A gradual increase in relative luminescence intensity (RLU) was observed in PMLECs treated with mFLuc/sLNP formulations (**Figure 2I**), ranging from the lowest-performing DDSEH to the top-performing HOSEH, with a few exceptions (e.g., DOSPeH). Notably, the best-performing group of sLNPs in vivo also showed strong transfection activity in PMLECs. Spearman correlation analysis confirmed a strong positive correlation between PMLEC transfection efficiency and total lung bioluminescence flux (r = 0.9333, p = 0.0007; **Figure 2J**). These results suggest that the ability to transfect primary lung endothelial cells in vitro is predictive of in vivo lung-targeted mRNA delivery. We next tested mFLuc delivery in primary human pulmonary arterial endothelial cells (HPAECs). A similar trend was observed: inactive sLNPs (e.g., DDSEH, DHSEOAc) produced negligible RLU, whereas active sLNPs (e.g., DOSPeH, HOSEH) yielded high RLU. Spearman analysis revealed a significant positive correlation between HPAEC transfection efficiency and total lung flux (r = 0.70, p = 0.0433). Overall, these findings indicate that both PMLEC and HPAEC models can potentially serve as predictive in vitro systems for evaluating sLNP-mediated mRNA delivery to lung endothelium, although further validation is necessary.

Lastly, we evaluated the relationship between sLNP-mediated mRNA accumulation in the lungs and their in vivo delivery performance. To measure mRNA accumulation, Cy5 -labeled FLuc mRNA (Cy5-mRNA) was encapsulated into different sLNP formulations and intravenously injected into mice. IVIS fluorescence imaging was then used to quantify Cy5 -mRNA distribution in the lungs. A clear trend emerged that the top-performing sLNPs produced stronger Cy5 epifluorescence signals, indicating higher mRNA accumulation in lung tissue, whereas low-performing sLNPs exhibited weak signals and limited accumulation (**Figure 2K**). Correlation analysis showed a strong positive correlation between lung Cy5-mRNA accumulation and delivery efficiency (r = 0.9667, p = 0.0002; **Figure 2L**). Collectively, these findings indicate that the most effective sLNPs not only promote enhanced mRNA accumulation in the lungs after systemic administration but also efficiently transfect pulmonary endothelial cells *in vitro*. Together, these properties likely underlie their superior lung-targeted mRNA delivery and protein expression *in vivo*.

#### Kinetics, subtype cell population identification, and proteomics analysis

To further characterize sLNP-mediated lung-targeted mRNA delivery, HOSEH, one of the top-performing sLNPs, was selected due to its high lung-targeting efficacy and specificity. The hydrodynamic diameter (<*D*_h_>) was determined to be 148.4 nm, with a PDI of 0.205, a zeta potential of 50.52 mV, and an encapsulation efficiency of 95.7%, all of which are comparable to our previously reported DOSEH and DHSEH sLNPs.^33^ The morphology was then examined by cryo-EM, revealing multilamellar vesicular structures as the primary morphology (**Figure 3A**). These structures are similar to the previously reported morphology of DOSEH, indicating that the asymmetric tail structure did not significantly affect sLNP self-assembly.^33^ Further analysis revealed a membrane thickness of approximately 2 nm, aligning well with the bilayer lipid membrane thickness (**Figure 3B**). Similar to previous findings, we speculate that the hydrophilic and negatively charged mRNA molecules interact with the positively charged bilayer membranes or are sandwiched between the multilamellar structures through electrostatic interactions and hydrogen bonding.^46, 47^

**Figure 3.**
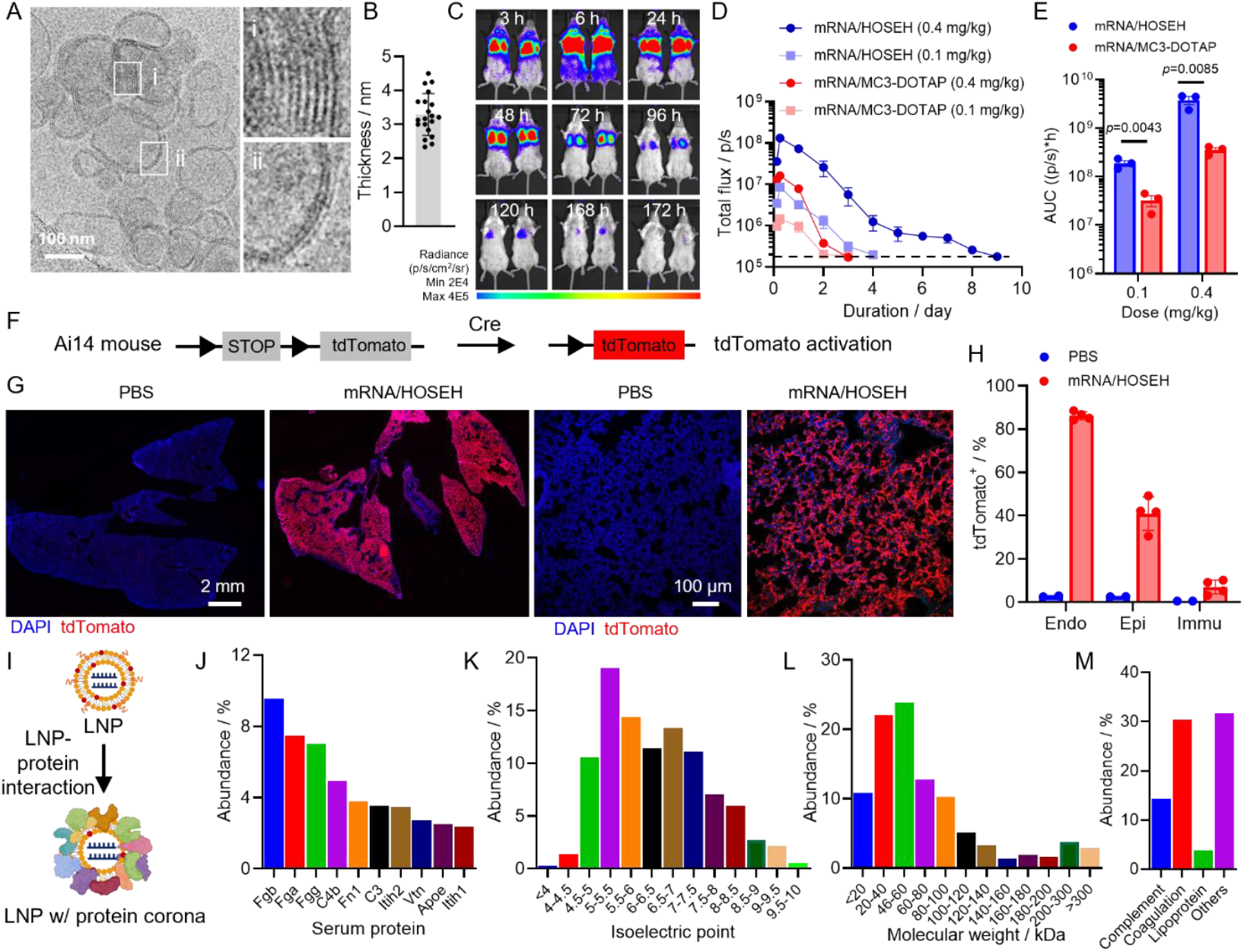
Characterization of mRNA/HOSEH formulation. (A) representative CryoEM images and (B) membrane thickness analysis of mRNA/HOSEH. (C) Time-dependent IVIS whole body bioluminescence images, (D) quantification of lung total flux, and (E) AUC analysis of mice treated with mRNA/LNP formulations. (F) Schematic illustration of Cre-mediated gene recombination in Ai14 mice. (G) Representative fluorescence images of lung lobes (left) and confocal images (right) and (H) flow cytometry analysis of tdTomato-positive lung cells of Ai14 mice treated with mCre/HOSEH. (I) schematic illustration of protein corona formation on sLNP following iv injection. Quantifications of (J) protein species, (K) PI, (L) molecular weight, and (M) functions of enriched proteins in the mRNA/sLNP corona. Data are presented as mean ± SEM, N = 3-4, student’s t-test.

Next, the protein expression kinetics of HOSEH sLNP-mediated mRNA delivery were studied. The lung-tropic MC3-DOTAP formulation was used as a control.^48-50^ Throughout this study, strong bioluminescence signals were observed primarily in the lung area for both formulations (**Figure 3C**). Additional signals in the auxiliary lymph node region were also observed in the mRNA/sLNP group. The mechanism underlying this observation remains unclear and merits further investigation. A dose-dependent total flux was observed for both the mRNA/sLNP and mRNA/MC3-DOTAP formulations, with higher doses inducing greater and longer expression (**Figure 3D**). In the mRNA/sLNP group, strong signals were detected at 3 hours post-injection, peaking at 6 hours and gradually decreasing thereafter. At a dose of 0.1 mg/kg, the bioluminescence signal returned to baseline by day 4 (96 hours), whereas a higher dose of 0.4 mg/kg produced a stronger intensity and prolonged expression, with signals persisting until 9 days post-injection (216 hours) before returning to baseline. Similarly, in the mRNA/MC3-DOTAP group, bioluminescence peaked at 6 hours post-injection and then gradually declined. At a dose of 0.4 mg/kg, signals returned to baseline by day 4 (96 hours), whereas at 0.1 mg/kg, signals diminished to baseline by day 2 (48 hours). Overall, at the same dose, the optimized HOSEH sLNP induced greater and more prolonged protein expression compared to the benchmark MC3 -DOTAP LNP. Furthermore, the area under the curve (AUC) was quantified as an indicator of total protein production following injection, showing that mRNA/sLNP led to significantly higher AUC at both low and high doses, highlighting the excellent efficacy of the sLNP delivery platform (**Figure 3E**).

In our previous study, lung cell population subtypes transfected by DOSEH and MC3 -DOTAP were quantified, identifying endothelial cells as the primary targets.^33^ In this study, we employed the same method by injecting Cre mRNA (mCre)-loaded HOSEH sLNP into adult Ai14 mice, which harbor a Cre-loxP reporter cassette in their genome (**Figure 3F**). Five days post-injection, tdTomato fluorescence signals were analyzed via fluorescence imaging and flow cytometry, using PBS-treated mice as controls. mCre/sLNP induced strong tdTomato fluorescence signals across all lung lobes, whereas no significant signal was observed in the PBS-treated control group (**Figure 3G** and **S6**). Flow cytometry analysis revealed that 80% of CD31^+^ endothelial cells, 50% of CD326^+^ epithelial cells, and 10% of CD45^+^ immune cells were tdTomato^+^ (**Figure 3H**).

Previous studies suggest that the protein corona may play a significant role in the organ-targeting mechanism following systemic administration (**Figure 3I**).^51-53^ The protein corona of HOSEH sLNP was isolated and analyzed via proteomics, revealing the association of over 200 different serum proteins, which is consistent with the DOSEH protein corona profile. The top enriched protein, accounting for 80% of the cumulative abundance, was further analyzed. The ten most abundant proteins included Fgb, Fga, Fgg, C4b, Fn1, C3, Itih2, Vtn, ApoE, and Itih1 (**Figure 3J**). These proteins differ from those found in the DOSEH corona, suggesting that hydrophobic tail structures influence serum protein adsorption.^33^ Approximately 80% of the proteins had a pI below 7.4, indicating a net negative charge at physiological pH, allowing complex formation with the positively charged sLNPs via electrostatic interactions (**Figure 3K**). Additionally, 60% of the proteins had molecular weights below 100 kDa, with the 20-40 kDa (22%) and 40-60 kDa (25%) ranges being the most abundant, similar to the DOSEH protein corona (**Figure 3L**). Proteins comprising 80% of the corona were categorized based on biological function: ∼12% were complement proteins, 30% were coagulation proteins, and 5% were lipoproteins (**Figure 3M**). Comparing HOSEH sLNP with other previously reported lung-targeting amine-based LNP systems, complement, coagulation, lipoproteins, and immunoglobulins were commonly present, reflecting the natural biological response to foreign particles. Previous reports suggested that vitronectin might mediate lung targeting through ligand-avβ3 interactions.^54, 55^ Consistent with this, vitronectin was identified as one of the most enriched proteins in the corona of HOSEH sLNPs. Additionally, alpha-1 antitrypsin (AAT), which was found enriched in the DOSEH protein corona, was also detected in the HOSEH corona.^33^

#### Safety and biocompatibility of sLNP

Beyond delivery efficacy and specificity, safety is a critical parameter for the clinical translation of mRNA/sLNP formulations.^56-58^ To evaluate the safety and biocompatibility of mRNA/sLNP in adult C57BL/6 mice, hematological toxicity was first evaluated, including total cell counts of red blood cells (RBCs), white blood cells (WBCs), and platelets (PLTs), as well as WBC differentials (lymphocytes, monocytes, and neutrophils). No significant differences were detected (**Figures 4A-F**), suggesting that systemic exposure to mRNA/sLNP was well tolerated without apparent hematotoxicity. Furthermore, potential hepatotoxicity and nephrotoxicity were assessed by measuring plasma levels of alanine transaminase (ALT), aspartate transaminase (AST), blood urine nitrogen (BUN), and creatinine (CRE). No significant differences were observed in these biochemical markers (**Figures 4G-J**), indicating that mRNA/sLNP did not induce liver or kidney toxicity.

**Figure 4.**
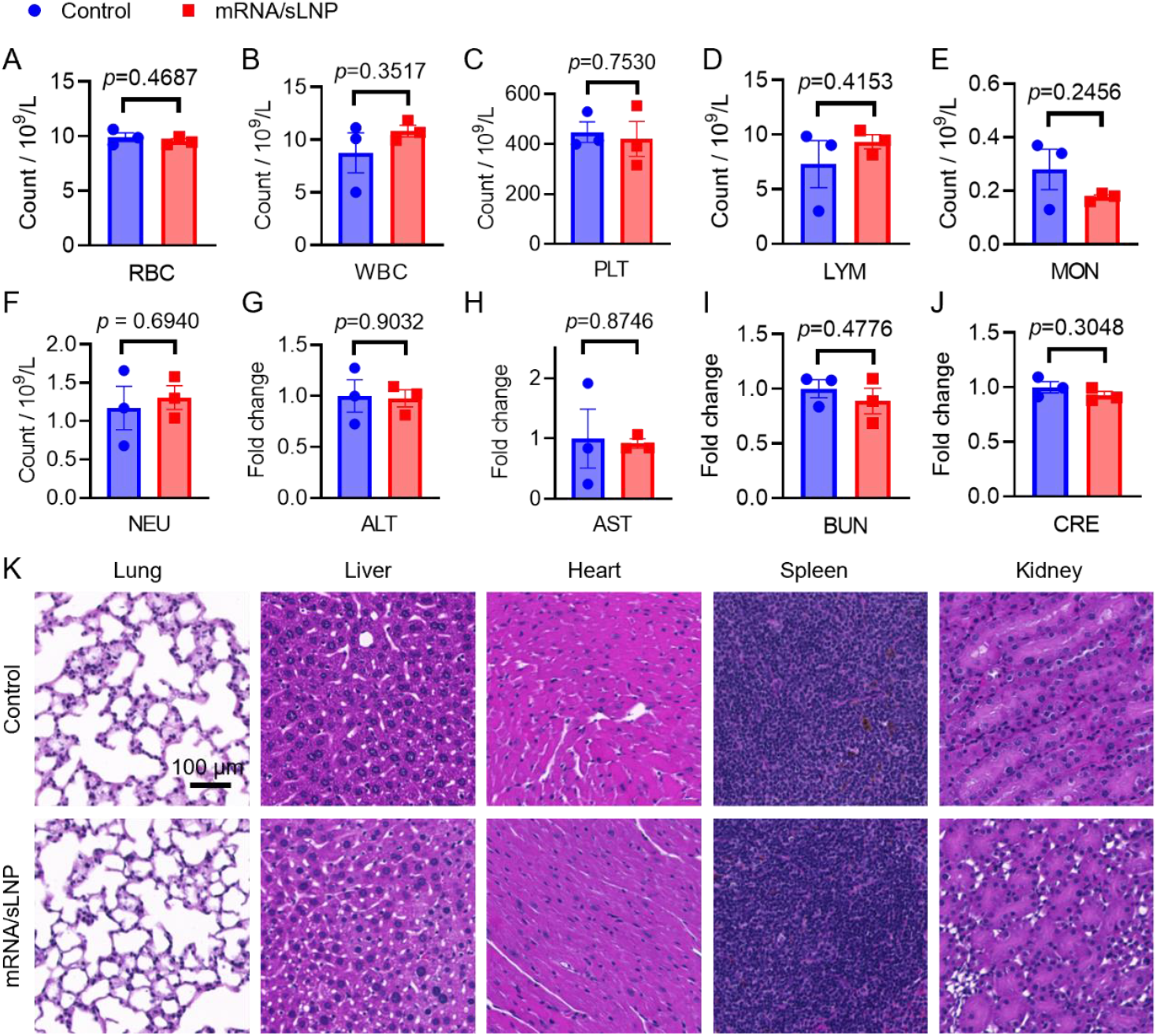
Toxicity study of mRNA/HOSEH formulation. Cell counts of (A) RBC, (B) WBC, (C) PLT, (D) LYM, (E) MON, and (F) NEU. Serum levels of (G) ALT, (H) AST, (I) BUN, and (J) CRE in mice treated with PBS or mRNA/sLNP. Data are presented as mean ± SEM, N = 3. (K) Representative H&E staining of tissue slices. schedule. mIL-10/sLNP was administered intravenously 2 h prior to LPS stimulation. (B) Western blot analysis and (C) quantification of IL-10 protein expression in lung tissues relative to control. (D, E) TNF-α and IL-6 levels in BALF, and (F, G) TNF-α and IL-6 levels in serum of mice treated with PBS or mIL-10/sLNP. (H, I) Neutrophil and macrophage counts in BALF. (J) Cytology analysis and (K) total protein concentrations in BALF of PBS- or mIL-10/sLNP-treated mice. Data are presented as mean ± SEM. N = 3.

Histological analysis of major organs, including the lung, liver, kidney, heart, and spleen, was performed using H&E staining (**Figure 4K**). Compared to PBS-treated control mice, no evident histopathological changes were observed in any examined organs, further demonstrating the safety of the sLNP formulation. Overall, the optimized HOSEH sLNP exhibited an excellent safety and biocompatibility profile in adult mice, providing a strong foundation for the future clinical translation of this lung-specific mRNA delivery system.

#### Prophylactic mIL-10/sLNP treatment ameliorates ALI in mice

Building on the efficacy and safety demonstrated by sLNP formulations with reporter mRNA, we next investigated the therapeutic potential of an immunomodulatory IL-10 mRNA-loaded HOSEH sLNP formulation (mIL-10/sLNP) in an LPS-induced ALI mouse model (**Figure 5A**).^38^ Following intravenous injection in healthy adult mice, IL-10 protein expression in lung tissue was first confirmed by western blot analysis (**Figure 5B**). Compared with mice treated with GFP mRNA-loaded sLNPs (mGFP/sLNP), those receiving mIL-10/sLNP showed significantly higher levels of IL-10 expression in the lungs (**Figure 5C**). To assess its prophylactic effects in ALI, mIL-10/sLNP was administered via intravenous injection 2 hours before intratracheal LPS stimulation (**Figure 5A**). Mice were sacrificed 24 hours post-LPS challenge, and bronchoalveolar lavage fluid (BALF) was collected for analysis. Compared to the untreated control group, LPS stimulation resulted in significantly elevated levels of proinflammatory cytokines TNF-α and IL-6 in BALF (**Figure 5D** and **E**). However, pre-treatment with mIL-10/sLNP significantly reduced TNF-α and IL-6 levels, suggesting that prior expression of IL-10 effectively suppresses inflammation in the lung environment (**Figure 5D** and **E**). Similarly, systemic TNF-α and IL-6 levels in serum were also markedly reduced in the pre-treatment group(**Figure 5F** and **G**), indicating that prophylactic mIL-10/sLNP administration mitigates both local and systemic proinflammatory cytokine production.

**Figure 5.**
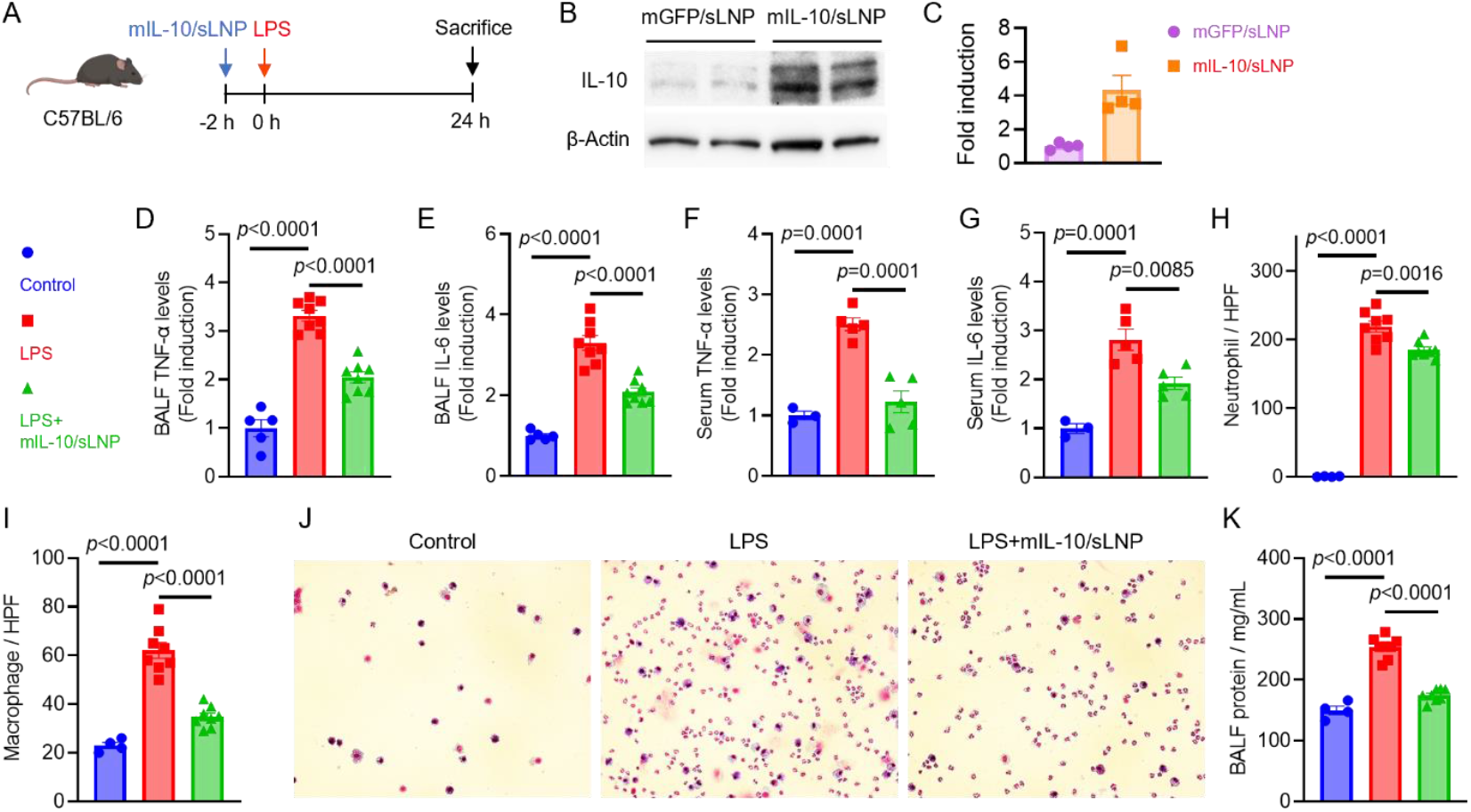
Prophylactic effects of mIL-10/sLNP in ALI mice. (A) Schematic of the treatment schedule. mIL-10/sLNP was administered intravenously 2 h prior to LPS stimulation. (B) Western blot analysis and (C) quantification of IL-10 protein expression in lung tissues relative to control. (D, E) TNF-α and IL-6 levels in BALF, and (F, G) TNF-α and IL-6 levels in serum of mice treated with PBS or mIL-10/sLNP. (H, I) Neutrophil and macrophage counts in BALF. (J) Cytology analysis and (K) total protein concentrations in BALF of PBS- or mIL-10/sLNP-treated mice. Data are presented as mean ± SEM. N = 3.

Cytological analysis of BALF further revealed that LPS stimulation led to a substantial infiltration of neutrophils and macrophages in the lung tissues (**Figure 5H** and **I**). In contrast, pre-treatment with mIL-10/sLNP significantly reduced the total number of neutrophils, macrophages, and overall cellular infiltration in BALF. The quantification of different cell populations confirmed the microscopic observations, demonstrating a significant reduction in both neutrophils and macrophages in BALF (**Figure 5J**). Furthermore, total protein concentration in BALF was measured as an indicator of capillary-alveolar barrier integrity (**Figure 5K**). LPS stimulation significantly increased total protein levels, suggesting endothelial damage and vascular leakage during lung inflammation. However, pre-treatment with mIL-10/sLNP effectively reduced total protein concentration, indicating partial restoration of the capillary-alveolar barrier, which is critical for maintaining lung health and function.

#### Therapeutic mIL-10/sLNP treatment ameliorates ALI in mice

To better reflect clinical scenarios in which patients typically present after inflammation has already developed, we evaluated the therapeutic potential of mIL-10/sLNP in attenuating LPS-induced ALI in mice^1, 3, 4^. Prior to testing IL-10 mRNA, we first examined how the inflammatory state of diseased lungs affects sLNP delivery and expression.^33, 35^ We began by examining the biodistribution of DiR-labeled mRNA/sLNPs in the lungs of ALI mice (**Figure 6A**). Epifluorescence imaging revealed no significant differences in lung fluorescence intensity between healthy and ALI groups, indicating that sLNPs accumulated in comparable amounts regardless of inflammatory status (**Figure 6B**). We next evaluated mRNA-encoded protein expression using FLuc mRNA-loaded sLNPs. In healthy mice, strong lung-restricted bioluminescence was observed within the 24-hour study period, showing a gradual increase from 3 to 6 hours, a peak at 6 hours, and a subsequent decline by 24 hours, consistent with the expected expression kinetics (**Figures 6C** and **D**). In ALI mice, robust lung-specific bioluminescence signals were also detected, confirming that sLNPs retained their targeting specificity under inflammatory conditions. However, the overall signal intensity in ALI lungs was significantly lower than that in healthy lungs, indicating reduced protein expression efficiency. This observation is consistent with prior reports that inflammatory conditions can impair cellular translation of delivered mRNA, and our results suggest that this principle also applies to the sLNP delivery platform.^59-61^ In one study, Lokugamage et al. reported that in a liver-targeting mRNA/LNP system, activation of TLR4 by LPS inhibited mRNA translation without affecting LNP uptake. Pharmacological inhibition of TLR4 or its downstream effector, protein kinase R (PKR), partially restored mRNA translation.^59^ It would be interesting to investigate the potential effects of pattern recognition receptor antagonists in lung-targeted delivery.

**Figure 6.**
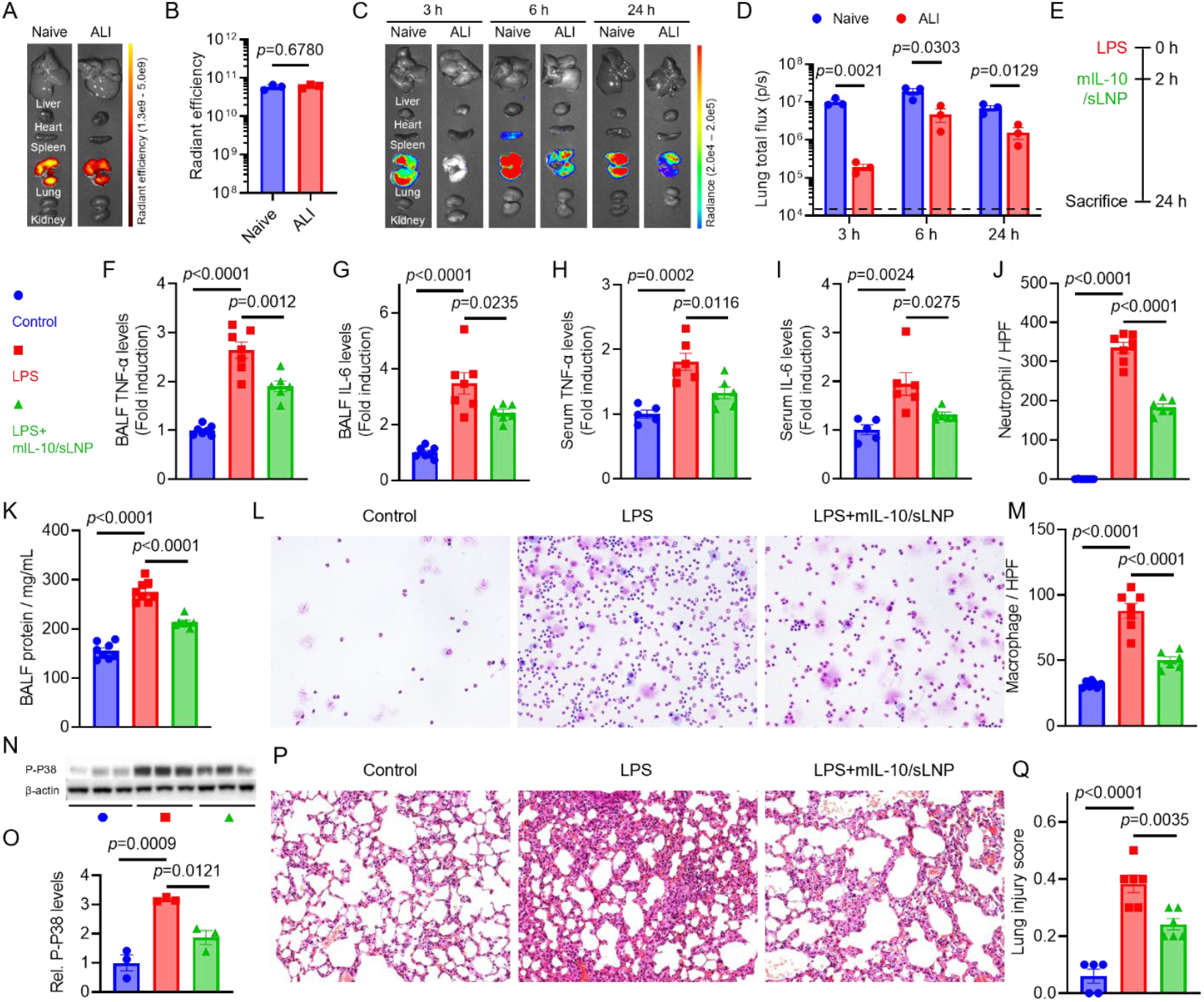
Therapeutic effects of mIL-10/sLNP in ALI mice. (A) IVIS epifluorescence images of organs from naïve or ALI mice treated with DiR-labeled mRNA/sLNP. (B) Radiant efficiency of lung tissues from naïve or ALI mice treated with DiR-labeled mRNA/sLNP. (C) Time-dependent IVIS bioluminescence images of ex vivo organs from naïveor ALI mice treated with mFLuc/sLNP. (D) Lung total flux of naïve or ALI mice treated with mFLuc/sLNP. (E) Schematic of the treatment schedule. mIL-10/sLNP was administered intravenously 2 h post LPS stimulation. (F, G) TNF-α and IL-6 levels in BALF, and (H, I) TNF-α and IL-6 levels in serum of PBS- or mIL-10/sLNP-treated mice. (J, K) Neutrophil and macrophage counts in BALF. (L) Cytology analysis and (M) total protein concentrations in BALF of PBS- or mIL-10/sLNP-treated mice. (N) Western blot analysis and (O) quantification of p-p38 expression in lung tissues. (P) H&E staining of lung tissues and (Q) quantification of lung injury scores in control, LPS, and LPS + mIL -10/sLNP groups. Data are presented as mean ± SEM. N = 3-7. Unpaired two-tailed Student’s t-test or one-way ANOVA.

Next, mIL-10/sLNP was administered via intravenous injection 2 hours after LPS stimulation. Mice were sacrificed 24 hours post-LPS challenge for analysis (**Figure 6E**). Consistent with the prophylactic study, the therapeutic administration of mIL-10/sLNP significantly reduced TNF-α and IL-6 levels in both BALF (**Figures 6F** and **G**) and serum (**Figures 6H** and **I**), confirming that mIL-10/sLNP effectively mitigates proinflammatory cytokine production. Cytological analysis further revealed that neutrophil and macrophage infiltration was markedly reduced in the treatment group compared to the LPS-only group (**Figures 6J, K** and **L**). Additionally, total protein concentration in BALF was significantly lower in the treatment group, indicating that mIL - 10/sLNP preserved endothelial-alveolar barrier integrity (**Figures 6M**). Collectively, these findings demonstrate that therapeutic mIL-10/sLNP treatment effectively reduces local and systemic inflammation, suppresses immune cell infiltration, and restores lung barrier function in ALI.

Previous studies have shown that IL-10 regulates proinflammatory cytokine production by inhibiting the activation of the p38 mitogen-activated protein kinase (MAPK) pathway.^20, 21^ To further investigate this mechanism, we analyzed phosphorylated p38 (p-p38) levels in lung tissues using Western blotting (**Figures 6N** and **O**). LPS-induced inflammation resulted in a significant increase in p-p38 expression compared to controls. However, mIL-10/sLNP treatment markedly reduced p-p38 levels, suggesting that inhibition of p38 activation may contribute to the therapeutic effects of IL-10 in LPS-induced lung inflammation. Next, we evaluated lung tissue damage in ALI mice and those treated with mIL-10/sLNP using H&E staining and lung injury scoring (**Figures 6P** and **Q**). LPS stimulation induced severe lung tissue injury, with an average injury score of 0.4, characterized by prominent neutrophil infiltration, alveolar hemorrhage, edema, and hyaline membrane formation. Treatment with mIL-10/sLNP significantly mitigated histopathological lung injury and decreased lung injury scores, highlighting its protective effects.

ALI/ARDS is associated with a spectrum of severe complications during the acute phase, among which acute kidney injury (AKI) is particularly common.^62-64^ We measured two established markers of kidney injury, serum creatinine and neutrophil gelatinase–associated lipocalin (NGAL). In the LPS-treated group, both markers were significantly elevated, indicating the development of AKI. In contrast, treatment with mIL-10/sLNP markedly reduced their levels, suggesting that IL-10 delivery not only mitigates ARDS pathology but also improves its associated complications.

Collectively, these findings demonstrate that although LPS-induced inflammation did not substantially alter mRNA/sLNP accumulation in ALI lung tissue, both mRNA delivery and subsequent protein expression were reduced within 24 hours. Nevertheless, robust lung-specific protein expression was still achieved. Importantly, administration of mIL-10/sLNP 2 hours after LPS stimulation effectively suppressed pulmonary and systemic proinflammatory cytokines, reduced immune cell infiltration, restored alveolar–capillary barrier integrity, and attenuated both lung injury and secondary renal damage.

#### Single-cell RNA sequencing

To further illustrate the transcriptional changes in individual lung cells induced by mIL-10/sLNP treatment, single-cell RNA sequencing was performed on lung tissues obtained following BALF lavage. Quality control procedures included filtering cells based on read counts, detected gene numbers, and mitochondrial content; identification and removal of doublets and ambiguous clusters; and evaluation of sample-level variability. It was found that, across all cells, LPS treatment was associated with enhanced expression of several genes implicated in inflammation (e.g., Tlr4, Nfkb1, Nfkb2, Tnf, Il1b), an effect that was mitigated in samples also treated with mIL - 10/sLNP (**Figure 7A**). To explore cell-type specific responses to LPS and LPS+mIL-10/sLNP, cluster analysis was performed to identify subsets of transcriptionally similar cells. In total, 16 clusters showed distinct transcriptional profiles that corresponded to those of 13 general cell types (**Figure 7B** and **C**). Transcriptional profiles of separate clusters were consistent with monocytes (23.9% of total cells), B cells (14.5%), endothelial cells (14.4%), neutrophils (11.6%), interstitial macrophages (10.7%), T cells and natural killer cells (10.3%), alveolar macrophages (4.0%), fibroblasts (3.6%), alveolar type I and type II cells (0.9% and 1.9%, respectively), dendritic cells (1.9%), mural cells (1.7%), and mesothelial cells (0.6%). **Figure 7D** and **E** present the number of cells from each sample in each cluster.

**Figure 7.**
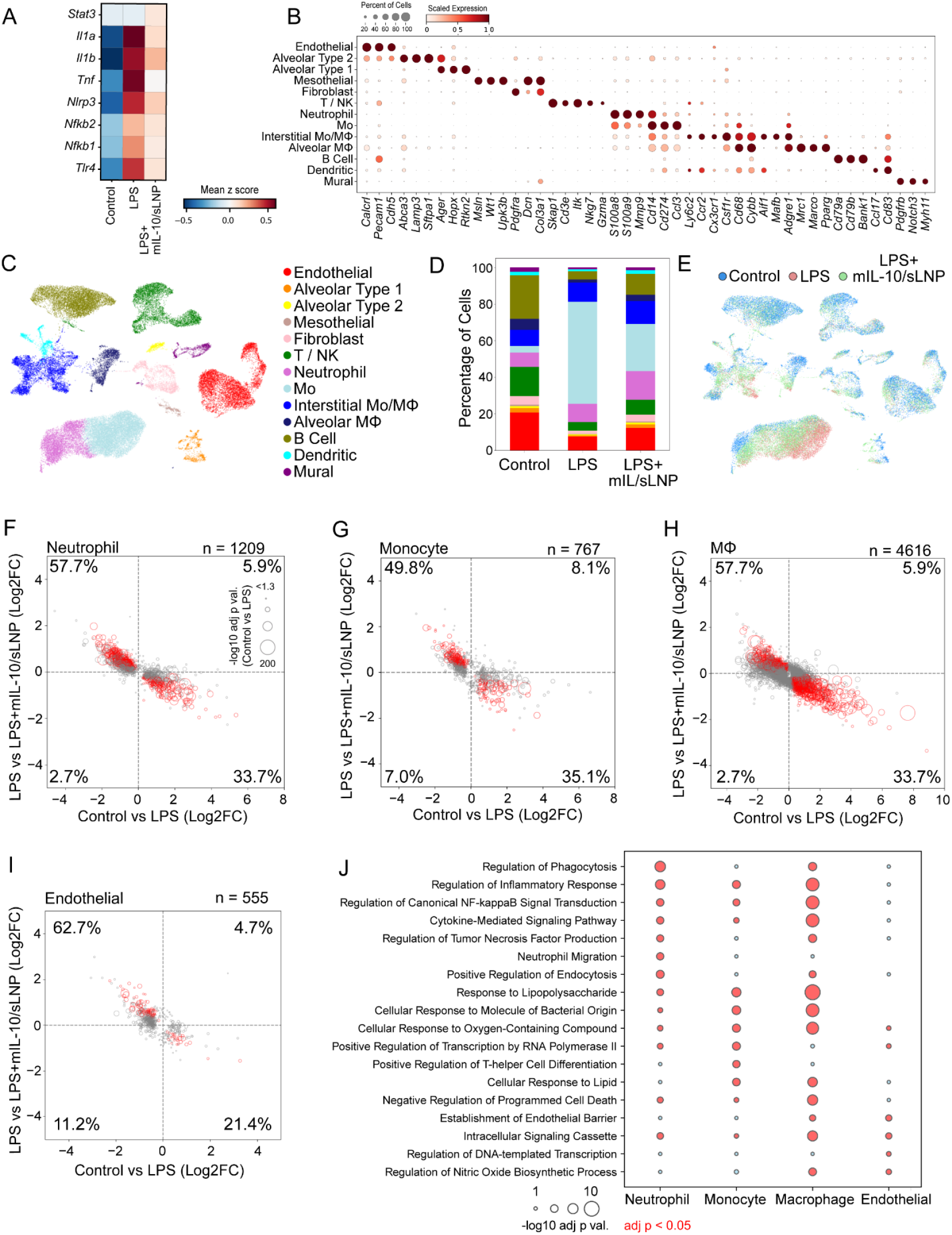
Single cell RNA sequencing reveals mitigation of LPS-induced transcriptional shifts by LPS+mIL-10/sLNP delivery. (A) Expression levels (mean z-score) across gene sets implicated in inflammatory pathways are presented across experimental groups, revealing a relatively large inflammatory signal in LPS-treated samples that is mitigated in LPS+mIL-10/sLNP samples. (B) Cluster analysis revealed 13 general cell classes largely showing selective expression of cell-type marker genes. The size and color intensity of each bubble correspond to the proportion of cells that show non-zero expression of a given gene and its level of expression scaled from 0-1 across the clusters, respectively. (C) UMAP of individual cells color coded by cell type shows clear separation of cell types in reduced dimensions. (D) The percent of cells within each cell-type is plotted across experimental groups. (E) UMAP of cells color coded by treatment group, (F-I) Scatter plots of log2 fold change values from LPS vs Control and LPS+mIL-10/sLNP vs LPS comparisons in (F) neutrophils, (G) monocytes (Mo), (H) macrophages (MΦ), and (I) endothelial cells show a largely anticorrelated profile across the two comparisons, indicating a ‘recovery’ profile when mIL-10/sLNP was delivered following LPS stimulation. The total number of genes included in the plot represent those showing significant (adj p < 0.05) differential expression in either comparison; those showing significant differential expression in both comparisons as well as an anti-correlation across comparisons are shown in red. The proportion of genes in each quadrant are presented. Positive values indicate upregulation in LPS (relative to Control) or LPS+mIL-10/sLNP (relative to LPS); negative values indicate upregulation in Control (relative to LPS) or LPS (relative to LPS+mIL-10/sLNP). (J) Subsets of functional terms (from the GO Biological Process 2025 database) associated with gene sets showing a ‘recovery’ profile are presented for each cell type. The size of each bubble is proportional to the adj p value; adj p values < 0.05 are shown in red; Mo = Monocyte, MΦ = Macrophage.

Genes differentially expressed across experimental conditions were further identified particularly in neutrophils, monocytes, macrophages, and endothelial cells. Relative to Control samples, LPS- stimulation induced significant transcriptional changes in 1,162 genes in neutrophils, 649 in monocytes, 3,857 in macrophages, and 467 in endothelial cells (**Figure 7F-I**). In neutrophils, monocytes, and macrophages, LPS-responsive differentially expressed genes (DEGs) were associated with cellular response to lipopolysaccharide and regulation of inflammatory response, among others, while cytokine signaling, particularly signaling by interleukins, tumor necrosis factor and NF-kB, was implicated in all four cell types, as were processes related to transcription and translation. Delivery of mIL-10/sLNP in LPS-stimulated mice counteracted many of the LPS-associated effects across the four cell types, as evidenced by largely anticorrelated changes in gene expression across the LPS (vs Control) and the LPS+mIL-10/sLNP (vs LPS) comparisons (**Figure 7F-I**). LPS+mIL-10/sLNP induced a significant shift of expression levels toward Control values in 35.3% of the LPS-responsive DEGs in neutrophils, 36,7% in monocytes, 31.4% in macrophages, and 22.5% in endothelial cells (**Figure 7F-I**), suggesting a “recovery” profile. Functional terms associated with gene sets showing this “recovery” profile were queried from the Gene Ontology Biological Process 2025 and Reactome 2024 databases, presented in **Figures 7J**, respectively. Findings show over-representation of numerous pathways that were also altered by LPS, for example, inflammatory processes, cytokine signaling, and transcription; ‘recovery’ gene sets showing an association with interleukin signaling are presented for each immune cell type. Taken together, these results indicate that the lung-targeting mIL-10/sLNP treatment broadly counteracts LPS-induced inflammatory transcriptional reprogramming across diverse lung cell types. Single-cell RNA-sequencing analysis revealed that LPS stimulation upregulated key inflammatory genes and pathways, including those involved in cytokine signaling, NF-κB activation, and interleukin-mediated responses, in multiple immune and structural cell populations. Treatment of mIL-10/sLNP substantially mitigated these effects, restoring a large fraction of differentially expressed genes toward baseline levels, particularly in neutrophils, monocytes, macrophages, and endothelial cells. Functional enrichment analyses confirmed that genes showing this “recovery” pattern were enriched in inflammatory and cytokine signaling pathways, underscoring the potent anti-inflammatory and homeostatic effects of mIL-10/sLNP at the single-cell level.

### Summary

In summary, building upon our previously developed HOSEH and DOSEH lipids,^33^ we examined how variations in both head and tail structures influence lung-targeted mRNA delivery in vivo. Several key structure–activity relationships were identified. First, lipids with linear alkyl chains of C16–C18 exhibited the highest efficacy compared with shorter or longer chains. Incorporating asymmetry further improved performance. Second, the hydroxyl group was essential, as caging it with an acetyl group greatly reduced activity. Substitution with an amide group did not significantly alter efficacy. Third, lipids with octadecyl tails and hydroxyethyl or hydroxypropyl head groups performed best. Extending the spacer between the sulfonium and hydroxyl groups gradually diminished delivery. Although these findings provide valuable insights, the precise mechanisms warrant further investigation.^46, 65^ Correlation analyses revealed that conventional physicochemical properties of sulfonium lipids and their sLNP formulations (cLogP, hydrodynamic diameter, PDI, zeta potential, and encapsulation efficiency) did not predict in vivo lung delivery. Instead, two factors showed strong positive correlations with lung-targeted efficacy: in vivo lung accumulation efficiency and in vitro transfection performance in both PMLECs and HPAECs. Kinetics study further demonstrated that sLNPs induced lung-specific protein expression in a dose-dependent manner, peaking at 6 hours post-intravenous injection and declining gradually thereafter. Endothelial, epithelial, and immune cells were successfully transfected following a single dose, though endothelial cells remained the primary targets, consistent with earlier findings from our group and others.^66-68^ Proteomics analysis of the protein corona revealed a complex composition of adsorbed proteins, which may provide mechanistic clues for lung targeting. ^54^

Safety assessments indicated that mRNA/sLNPs were well tolerated in mice, with no evidence of hematologic, hepatic, or renal toxicity by biochemical analysis and no significant tissue damage. However, clinically relevant dosing levels have not yet been established for sLNP formulations, and dose-escalation studies will be necessary to identify potential dose-related toxicities. Finally, in an LPS-induced ALI mouse model, mIL-10/sLNP demonstrated both prophylactic and therapeutic efficacy. LPS-induced inflammatory transcriptional changes across multiple immune cell types in the lung, restoring gene expression toward normal levels. Despite these encouraging findings, several important challenges remain. In clinical practice, patients present at varying stages of disease progression, underscoring the need to optimize both therapeutic dosing and timing.^1-4^ In addition, ALI arises from diverse etiologies, including infection and sepsis, and the efficacy of mIL-10/sLNPs across these contexts has yet to be established. ^3, 4^ Notably, our sLNP platform has also demonstrated successful delivery of small-molecule drugs and antibiotics, suggesting that hybrid therapeutic strategies may provide a viable approach to address this complexity.^37, 38^

## Author Contribution

Y.M., R.C., Q.M., and Y.L. conceived and designed this study. Y.M., D.P., Y.S., Z.C., R.G., R.K., C.W., N.T., R.C., Q.M., and Y.L. performed experiments and analyzed data. All authors discussed and commented on the results. Y.M. and Y.L. wrote the manuscript. All authors read, commented on, revised, and approved the manuscript. Q.M. and Y.L. supervised the project.

## Acknowledgments

We thank Drs. Juntao Luo, Hong Lu and Changying Shi for their assistance with material characterization, Drs. Jennifer Moffat for bioluminescence imaging, Dr. Adam Waickman and Lisa Phelps for flow cytometry analysis, Dr. Jushuo Wang for confocal imaging, and Dr. Stephan Wilkens for CryoEM imaging. CryoEM images were collected in the Electron Microscopy Core facilities at SUNY Upstate Medical University and SUNY Environmental Science and Forestry (ESF). We thank the staff members at SUNY ESF Analytical and Technical Services, SUNY Upstate Medical University Research Flow Core, Pathology Research Core, and the Department of Laboratory Animal Resources. Some elements in Figures 1, 3, and 5 were created by BioRender.com. Y. L. acknowledges the Francis Hendricks Endowment Fund, NIH grants EB032579 and GM160083, and the startup fund from SUNY Upstate Medical University.

## Declaration of AI and AI-assisted technologies in the writing process

During the preparation of this work, the authors used ChatGPT for spelling checks, grammar checks, and rephrasing. After using this tool, the authors reviewed and edited the content as needed and take full responsibility for the content of the publication.

## Data Availability

The data that support the findings of this study are available from the corresponding author upon reasonable request.

